# Conservation genetics of the white-bellied pangolin in West Africa: a story of lineage admixture, declining demography and wide sourcing by urban bushmeat markets

**DOI:** 10.1101/2023.03.09.531886

**Authors:** Koffi Jules Gossé, Sery Gonedelé-Bi, Sylvain Dufour, Emmanuel Danquah, Philippe Gaubert

## Abstract

During the last 40 years, the volumes of African pangolins feeding the illegal wildlife trade have dramatically increased. We conducted a conservation genetics survey of the most traded African species, the white bellied pangolin (WBP; *Phataginus tricuspis*), across three West African countries including Guinea, Côte d’Ivoire and Ghana. Our study combining mitochondrial DNA (mtDNA) sequencing and microsatellites genotyping is the first to reveal a global pattern of admixture between two of the six mitochondrial lineages as previously delimited within WBP. We found a signature of isolation-by-distance and a lack of population genetic structuring, supporting the idea that WBP may have underestimated dispersal abilities. Levels of genetic diversity were low compared to central African lineages, reinforcing the picture of genetic pauperization shared by West African WBP. We observed a 85-98% decline in the effective population size of WBP occurring c. 3200 to 400 ya, with current numbers (520–590) at the lower end of the conservative thresholds for minimum viable population size. The microsatellites markers were powerful enough to differentiate between individuals and identify replicated samples, confirming the utility of this approach in tracing the pangolin trade. Genetic diversity estimates confirmed that Yopougon, the main bushmeat market from Abidjan (Côte d’Ivoire), was fed by a large trade network as confirmed by vendors reporting 10 different sources situated 62-459 km away from the market. We conclude that WBP distributed in the Upper Guinean Block should be considered a single management unit of high conservation concern, as impacted by genetic diversity erosion, drastic decline in effective population size and wide range sourcing for feeding urban bushmeat markets. Given the genetic admixture pattern detected within WBP from West Africa, we advocate for a multi-locus strategy to trace the international trade of the species.

## Introduction

Pangolins (Mammalia, Pholidota) are considered the most trafficked mammals in the world (Challender, Harrop & MacMillan, 2015; Heinrich *et al*., 2017; Challender *et al*., 2020), with approximately 900,000 individuals seized in the last 20 years (Challender *et al*., 2020). The eight species of African and Asian pangolins are threatened with extinction, through the cumulative effect of illegal wildlife trade and deforestation (Heighton & Gaubert, 2021). As a consequence, they all have been listed on Appendix I of the Convention on International Trade in Endangered Species of Wild Fauna and Flora (CITES) (Challender & Waterman, 2017), and rated as Vulnerable, Endangered or Critically Endangered on the IUCN Red List of Threatened™ Species.

In tropical Africa, land conversion, deforestation and hunting are the major drivers of faunal depletion, including pangolins (Pietersen, McKechnie & Jansen, 2014a; Boakye *et al*., 2016; Ingram *et al*., 2018). Pangolins have long been hunted across their ranges as part of the traditional game spectrum consumed by local communities (Boakye *et al*., 2014; Zanvo *et al*., 2021). However, the large demand from the Chinese traditional medicine (Cheng, Xing, & Bonebrake, 2017; Challender & Waterman, 2017) and the apparent decrease in numbers of Asian pangolins (Challender & Waterman, 2017; Heinrich *et al*., 2017), have recently created a global source-sink trade network where pangolin scales (mostly) are being massively exported from Africa to South-East Asia (Challender & Hywood, 2012; Ingram *et al*., 2019; Zhang *et al*., 2020).

From 1972 to 2014, the harvest of African pangolins has dramatically increased with volumes being multiplied by nine between 2005 and 2014 (Ingram, Coad & Scharlemann, 2016). For central Africa alone, the amount of annually hunted pangolins has been estimated to increase from 0.42 to 2.71 M animals (Ingram *et al*., 2018). Despite being –wrongly– blamed for their role as intermediate hosts of the COVID-19 pandemic (Frutos *et al*., 2020), pangolins are still harvested at high rates (Aditya *et al*., 2021), with an estimate of > 400,000 African pangolins bound for Asian markets between 2015 and 2019 (Challender *et al*., 2020).

In Côte d’Ivoire (CI), the bushmeat trade remains a vibrant activity despite the national hunting ban established in 1976 (Caspary, Koné, & De Pauw, 2001; Gonedelé Bi *et al*., 2012). Three species of pangolins can be found on the bushmeat stalls, including the white-bellied pangolin (*Phataginus tricuspis*), the black-bellied pangolin (*P. tetradactyla*) and the giant pangolin (*Smutsia gigantea*), although the latter is now subject to local extirpation (Gonedelé Bi *et al*., 2017). There is growing evidence that part of the pangolin trade in CI is now feedin an non-domestic, international network, as testified by seizures of several hundreds of scales bound to South-East Asia and China during the last 15 years (Challender & Hywood, 2011; Ingram *et al*., 2019; Challender *et al*., 2020). Recently, Abidjan, the largest city in CI, was highlighted as one of the major western African hubs of the pangolin trade, with a record seizure of >3.5 tons of scales representing c. 10,000-15,000 pangolins (https://www.20minutes.fr/monde/2732399-20200304-cote-ivoire-saisie-35-tonnes-ecailles-pangolin-incineree-autorites).

Although the white-bellied pangolin (WBP) is the most trafficked African species (Zhang *et al*., 2020), its natural history remains poorly known, and as a correlate, so remains the genuine impact of the trafficking activities on its populations. Gaubert *et al*. (2016), described six geographically traceable mitochondrial lineages within WBP, one of which (West Africa; WAfr) being found in Côte d’Ivoire. Such lineage delineation was recently applied to the tracing of the global pangolin trade (Zhang *et al*., 2020). However, fragmentary knowledge on the lineages’ precise ranges –notably relative to the neighboring Ghana (Gha) lineage– and the lack of fine-scale resolution of mitochondrial DNA (mtDNA) markers, prevent from addressing the conservation genetics of WBP at a local scale.

Here, we investigate the genetic status of the species across two West African neighbor lineages (WAfr and Gha), building on the assumption that the genetic toolkit is a useful contributor to conservation-oriented implementations such as population-based management of pangolins (Zanvo *et al*., 2022). We set up an approach combining mtDNA to species-specific, recently developed microsatellites markers (Aguillon *et al*., 2020), that will contribute to filling the population genetics’ knowledge gap that the species remains subject to, notably in West Africa (Heighton & Gaubert, 2021). Our specific objectives were to assess (i) the genetic structure of WBP in West Africa and (ii) their genetic diversity and historical demography. Based on these outputs and an important sampling effort in several major bushmeat selling sites, we eventually discuss the potential of the genetic toolkit to contribute to the tracing of the pangolin trade in the subregion.

## Material and Methods

### Sample collection and wet laboratory procedure

Our study was conducted under research authorization 0632/MINEF/DGFF/FRC-aska issued by the Direction générale des forêts et de la faune du Ministère des Eaux et Forêts, Côte d’Ivoire. After explaining the objectives of the study, free consent was obtained from market and restaurant vendors prior to sample collection, without providing financial incentives. We collected a total of 116 samples from bushmeat markets, restaurants and food stalls across West Africa, including CI (102 samples; eight localities), Ghana (five samples; three localities) and Guinea (nine samples; one main area) (Appendix Table 1); so that sampling coverage captures the West Africa (WAfr) and Ghana (Gha) mitochondrial lineages’ ranges (Gaubert *et al*., 2016). The sample scheme was designed so to include both (i) reference samples that will serve for the genetic assessment of WBP and (ii) samples from main bushmeat markets (i.e., collected from large urban markets) for assessing trade traceability in the subregion (Fig. 1). Reference samples were defined as collected from selling places (rural areas outside of the major urban markets in the city of Abidjan) that sourced pangolins within a <40 km radius (there was no radius overlap among sites). The radius was delimitated through questionnaires addressed to 6-29 sellers from each site. Because our study scale is large and covers c. 1200 km, we consider that such selling places can be used as proxy of ‘populations’, as long as we could delimitate the proximity of their sourcing localities. Samples consisted of fresh or smoked tissues (tongue and muscle), which were preserved in 95% ethanol and stored at 4°C until laboratory processing.

**Fig 1.**
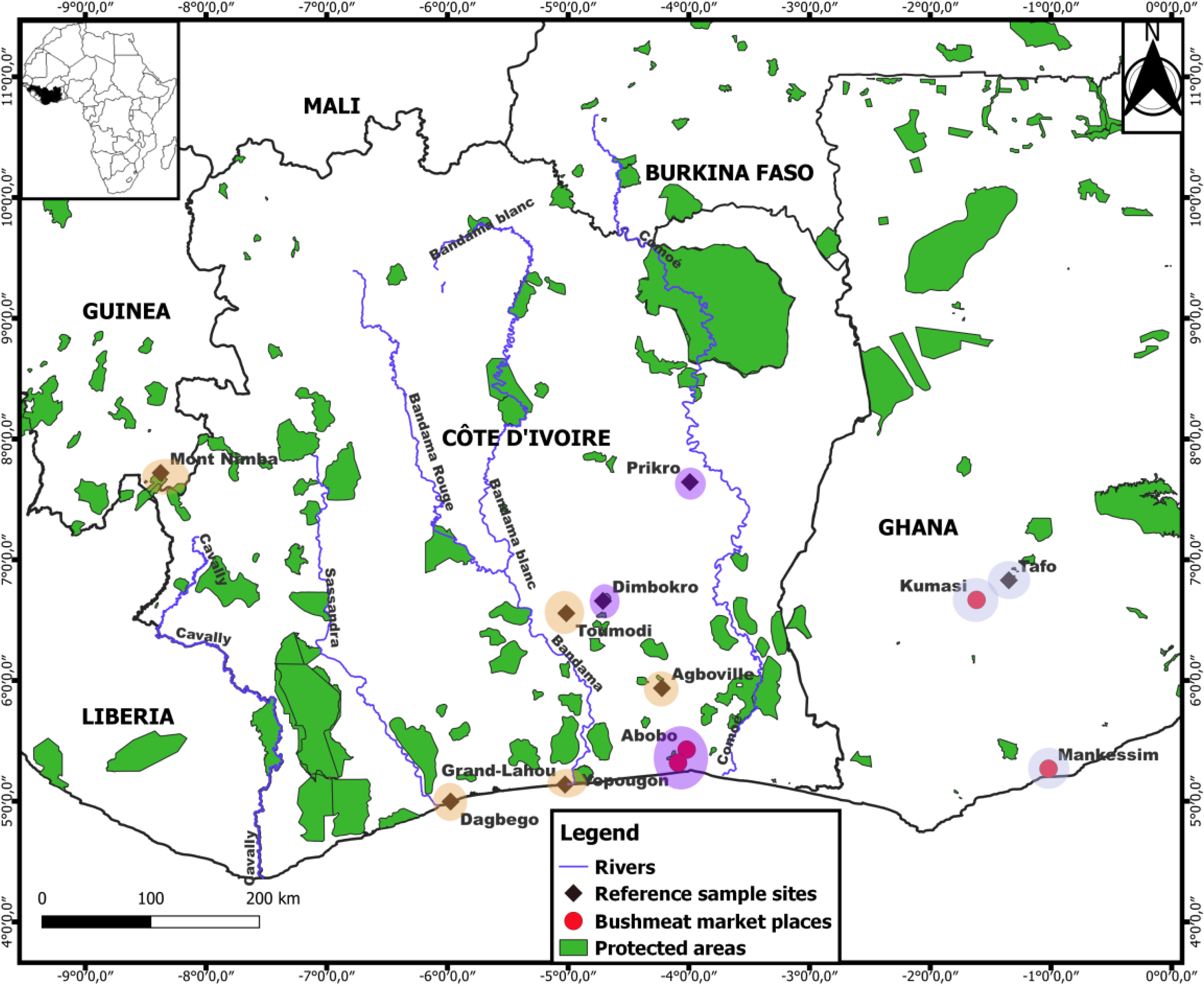
Re-assessed distribution of the Western Africa and Ghana mitochondrial lineages of white-bellied pangolins across West Africa. Blue and orange circles correspond to sampling sites where the Ghana (east) and Western Africa (west) mitochondrial lineages occur, respectively. Purple circles mean that both lineages co-occur. Bushmeat market samples originate from large urban markets and have no precise origin. Reference samples come from circumscribed sites (see Material and Methods).

DNA extraction was performed using the NucleoSpin® Tissue kit (Macherey-Nagel, Hoerdt, France), following manufacturer’s recommendations, with 50 μl final elution step repeated twice in order to increase DNA yield. DNA concentrations were estimated with the NanoDrop 1000 spectrophotometer (ThermoFisher Scientific, Illkirch Graffenstaden, France).

We amplified an mtDNA fragment of 432 bp from the control region (CR1), following Gaubert *et al*. (2016). PCR products were visualized on 1.5% agarose gel and sent for bidirectional sequencing at Genoscreen (https://www.genoscreen.fr/en; Lille, France) and Macrogen Europe (https://dna.macrogen-europe.com/en; Amsterdam, The Netherlands). Sequences were aligned manually with BioEdit v7.2.5 (Hall, 1999), and deposited in Genbank under accession numbers OP897333 - OP897461.

We amplified 20 microsatellite markers developed from the genome of WBP using four PCR multiplexes after Aguillon et al. (2020). Six loci (PT_276641, PT_353755, PT_378852, PT_1162028, PT_1753627 and PT_1849728) from the original multiplexes were not considered in our analysis, because of a significant proportion of amplification failures likely due to fluorescent dye degradation (for each locus, > 43% of missing data across all individuals). Serial PCR triplicates were randomly performed to mitigate potential issues related to allele dropout and null alleles in the 14 remaining loci. Consensus was considered achieved when at least two of the three replicates indicated the presence of an allele (Dayon *et al*., 2020). The PCR products were separated via capillary electrophoresis at Genoscreen.

### Genetic analyses – Control Region

#### Phylogenetic clustering

We evaluated the clustering of our CR1 sequences (N = 108) from West Africa relative to the six WBP mtDNA lineages (Gaubert *et al*., 2016) through a distance tree analysis including all the CR1 sequences available in Genbank at the time of drafting the manuscript (N = 101). Phylogenetic tree reconstruction was performed in MEGA 11 (Tamura, Stecher & Kumar, 2021) using Neighbor-Joining (NJ), 500 bootstrap replicates and Kimura 2-parameter model (Kimura, 1980).

#### Genetic diversity and structure

Genetic diversity and structure within WAfr and Gha were reassessed and compared to previous estimates (Zanvo *et al*., 2022). We used DnaSP v 6.12.03 (Rozas *et al*., 2017) to calculate for each lineage the number of haplotypes (h), mean haplotype diversity (Hd) and mean nucleotide diversity (π). We used Network as implemented in POPART v 1.7 (Leigh & Bryant, 2015) to build a haplotype median-joining network, with ε = 0 to minimize the number of alternative median junctions. We subsequently mapped haplotype distribution of the two lineages in QGIS v 3.22.2 (https://changelog.qgis.org/en/qgis/version/3.22/).

### Genetic analyses - Microsatellites

#### Genetic diversity

Geneious v 9.0.5 (Kearse *et al*., 2012) and the Microsatellites plugin (https://www.geneious.com/features/microsatellite-genotyping/) were used for allele scoring and genotype extraction. After excluding the six deficient loci (see above), we obtained a final dataset of 116 samples genotyped at 14 loci with at most 21% missing data per genotype (Appendix Table 1).

Validation of our 14 selected microsatellite loci was performed on a subset of “best” 24 samples with ≥80% genotyping success, taken from the whole study zone (as preliminary results showed a lack of nuclear genetic structure between WAfr and Gha; see below). Detection for the presence of null alleles, assuming population at equilibrium, was performed with Microcheker v 2.2.3 (Peakall & Smouse, 2012). We assessed deviation from Hardy-Weinberg equilibrium for each locus in GenAlEx v 6.503. We performed a permutation test in Arlequin 3.5 (Excoffier & Lischer, 2010) to estimate linkage disequilibrium (LD) between each pair of loci with 1,000 randomizations. Bonferroni correction was applied in the three above-mentioned analyses.

Genetic diversity for each *a priori* population with N ≥ 5 (except in Dagbégo: N = 3), including both reference populations (Dagbégo, Dimbokro, Toumodi, Prikro in Côte d’Ivoire, and the area of Mont-Nimba in Guinea) and market populations (Yopougon and Abobo in Abidjan, Côte d’Ivoire), was estimated through the number of alleles (Na) and private alleles (pA), observed (Ho), expected (He) and unbiased expected (uHe) heterozygosity (in GenAlEx), allelic richness (A_R_) and Wright’s fixation index (F_IS_) (in FSTAT v 2.9.4) (Goudet, 2003). We used the effective number of alleles (Ne) (Brown & Weir, 1983), as calculated from the reference populations, as an estimate of genetic diversity to compare among WBP lineages.

#### Discriminative power of microsatellites markers

We used GenAlEx to evaluate the discriminative power of our microsatellite markers in our total dataset (N = 116) by calculating the number of identical genotypes among samples with the *Multi-locus tagging* option (*Matches* sub-option). We used the *psex()* function from the *R* package *poppr* (with method = multiple; Kamvar *et al*., 2014) to assess the probability of encountering the same genotype more than once by chance. We ran Gimlet v1.3.3 (Valière, 2002) to calculate values of unbiased identity probability (uPI) and sibling identity probability (PIsibs).

#### Population structure

We used GenAlEx to perform a principal coordinate analysis (PCoA) with unbiased pairwise genetic distances in order to explore genetic variance among all WBP individuals from West Africa.

We conducted a clustering analysis in STRUCTURE v 2.3.4 (Pritchard, Wen, & Falush, 2010) including the 116 genotyped individuals. We performed 20 independent simulations with K values ranging from 2 to 8, using 10^5^ Markov chain Monte Carlo (MCMC) iterations and burnin = 10^4^, assuming admixture model and correlated allele frequencies. We used STRUCTURE HARVESTER v 0.6.94 (http://taylor0.biology.ucla.edu/structureHarvester/) to assess the most likely number of populations (K) according to the method of Evanno *et al*. (2005). The results of the 20 iterations for each K value were summarized and averaged with CLUMPAK (http://clumpak.tau.ac.il/contact.html) (Kopelman *et al*., 2015).

Pairwise differentiation between reference populations was estimated by calculating the fixation index (F_ST_) (Nei, 1986) with a randomization process of 1000 permutations to obtain p-values in Arlequin v 3.5 (Excoffier & Lischer, 2010).

We tested isolation by distance (IBD) (Bohonak, 2002) by running a Mantel test with the *adegenet* package in RStudio v 4.0.5 (R Development Core Team 2021), where we quantified the correlation (*r*) between genetic (Edward’s) and geographic (Euclidean) distances among individuals and populations.

#### Demographic history

Given the lack of structure observed between WAfr and Gha (see Results), we retraced the demographic history of West African WBP as a unique nuclear population using the *R* package *varEff* (Nikolic & Chevalet, 2014), an approximate likelihood method that infers temporal changes in effective population size. We followed Zanvo *et al*. (2022) in fixing a generation time of two years and average mutation rate of 5.10^-4^ per site per Myr. We ran the single mutation, geometric mutation and two-phase mutation models, using 10,000 MCMC batches with a length of 1 thinning every 100 lots and JMAX = 3. Proxies of confidence intervals for ancestral and current effective population size estimates were calculated from the harmonic mean distribution of each mutation model.

## Results

### Control region

Our NJ tree based on 209 mtDNA sequences recovered the six WBP geographic lineages (Gaubert et al., 2016) with robust node supports (> 80%), including Western Africa (WAfr), Ghana (Gha), Dahomey Gap (DG), Western Central Africa (WCA), Gabon (Gab) and Central Africa (CA) (Appendix Fig. 1). All the newly produced sequences from Guinea (N = 6) and Ghana (N = 2) clustered within WAfr and Gha, respectively. The 103 new sequences that we generated from CI clustered both within WAfr (Agboville, Abobo, Dagbégo, Dimbokro, Grand-Lahou, Prikro, Toumodi and Yopougon; N=77) and Gha (Abobo, Dimbokro, Prikro and Yopougon; N=26).

Among WBP mitochondrial lineages, levels of genetic diversity (Pi) were the highest for CA and WCA, and the lowest for West African lineages, WAfr showing the second lowest Pi value after DG (Table 1). In total, 28 CR1 haplotypes were identified across WAfr and Gha (Appendix Table 2). The haplotype network showed that WAfr and Gha were separated by 7 mutations, whereas 1-2 mutations separated within-lineage haplotypes (Fig. 2). We did not observe any clear geographic structuring of haplotypes within each lineage. Gha extended into the eastern territory of Côte d’Ivoire (Prikro and Dimbokro; Appendix Fig. 2). Hap2 was dominant in WAfr (found in 24 samples) and had a large distribution, from SE Guinea to eastern Côte d’Ivoire. Eleven haplotypes were found in the Yopougon bushmeat market, whereas only three haplotypes were unique to the reference populations. Nine Gha haplotypes were found in Yopougon and Abobo markets (Appendix Table 2).

**Table 1.** Mitochondrial diversity among white-bellied pangolin lineages. Lineages re-assessed as part of this study appear in bold. N: number of sequences; S: number of polymorphic sites; h: number of haplotypes; Hd: haplotype diversity; Pi: nucleotide diversity. SD = standard deviation. Gab was not considered as represented by a single sample.

| Lineages | N | S | h | Hd | Pi |
| --- | --- | --- | --- | --- | --- |
| | | | | Mean ± SD | Mean ± SD |
| **Western Africa (WAfr)** | **103** | **15** | **19** | **0,89 ± 0,019** | **0,00545 ± 0,00034** |
| **Ghana (Gha)** | **41** | **9** | **12** | **0,913 ± 0,018** | **0,00628 ± 0,00036** |
| Dahomey Gap (DG) | 13 | 8 | 9 | 0,936 ± 0,051 | 0,00516 ± 0,00066 |
| Western Central Africa (WCA) | 61 | 25 | 32 | 0,958 ± 0,012 | 0,00745 ± 0,00056 |
| Central Africa (CA) | 11 | 10 | 8 | 0,891 ± 0,092 | 0,00825 ± 0,00108 |

**Fig 2.**
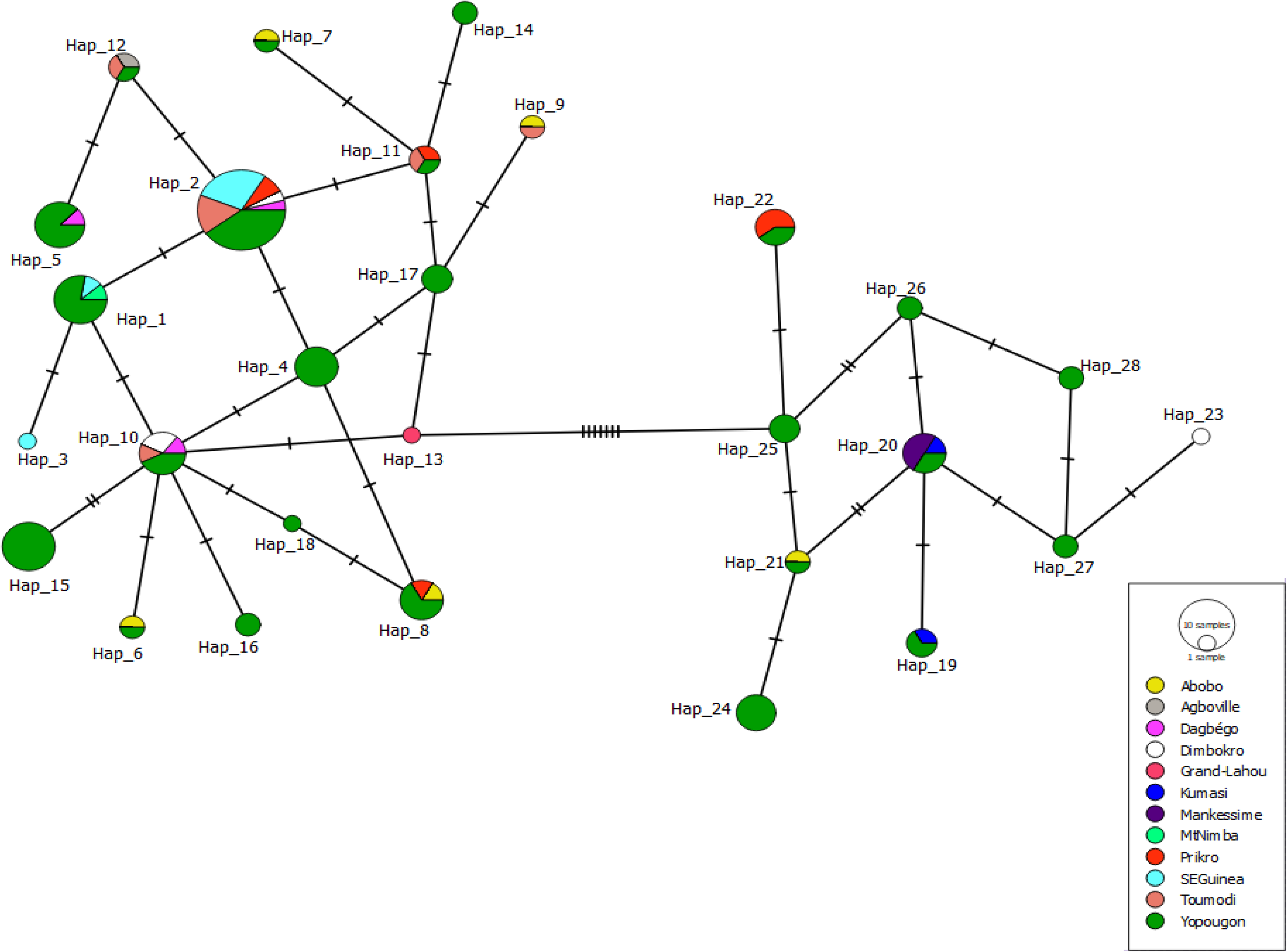
Median joining network of control region (CR1) haplotypes across the Western Africa and Ghana mitochondrial lineages of white-bellied pangolins. Haplotype numbers (e.g., Hap_8) refer to Appendix Table 2. Small perpendicular bars represent mutations.

### Microsatellites

Five samples shared the same genotype including (i) Yop68, Yop72 and Yop73 and (ii) Yop90 and Yop98. The null hypothesis of encountering the same genotypes more than once by chance was rejected for all pairs of individuals (P < 0.0001). We retained only Yop68 and Yop90 from each respective batch for downstream analysis.

Two loci significantly deviated (P < 0.003) from Hardy-Weinberg equilibrium (HWE) (Appendix Table 3). No pairs of loci were involved in LD. Null alleles were identified at three loci, involving the two loci deviating from HWE (Appendix Table 4). Mean number of alleles (Na) ranged from 3.000 (Dagbégo) to 8.928 (Yopougon), with mean value across populations = 4.673 (Table 2). Observed heterozygosity (Ho) and expected heterozygosity (He) ranged from 0.572 to 0.700 (mean = 0.648) and 0.549 to 0.652 (mean = 0.587), respectively. The values of uHe were similar among populations and market places, and ranged from 0.637 to 0.69 (mean = 0.656). F_IS_ values were significantly positive for the Dagbego and Mont-Nimba populations (0.009 and 0.067, respectively) and Toumodi (0.103). The number of private alleles ranged from 0 to 6 (Mont Nimba) in reference populations, and was 1 and 45 in Abobo and Yopougon markets, respectively. Mean effective number of alleles (Ne) across reference populations was 3.099.

**Table 2.**
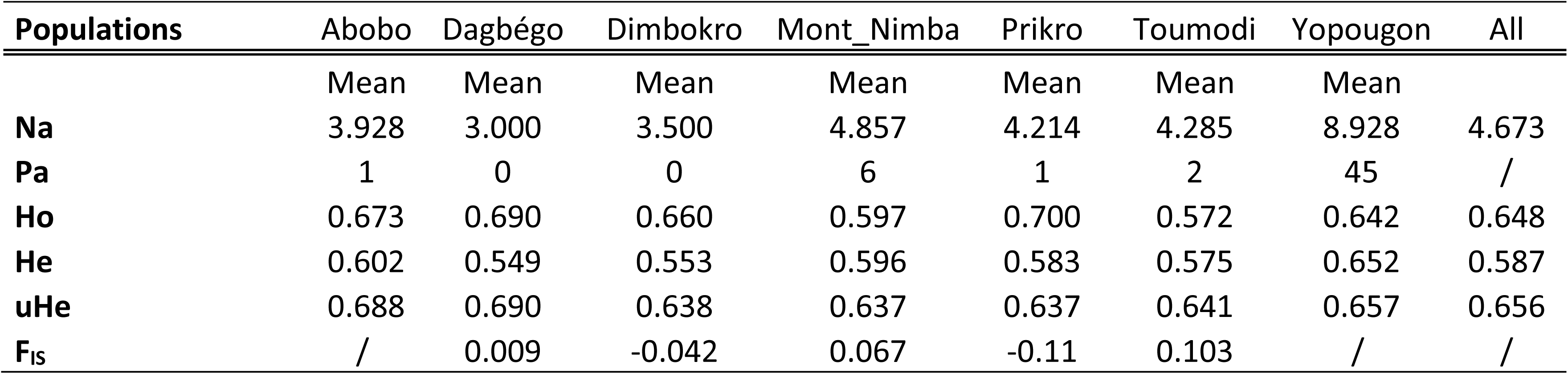
Diversity indices estimated from 14 microsatellite markers in populations of white-bellied pangolins from West Africa. Na: mean number of alleles; Pa: mean number of private alleles; Ho: observed heterozygosity; He: expected heterozygosity; uHe: unbiased expected heterozygosity; F_IS_: fixation index. Abobo and Yopougon are urban bushmeat markets.

As calculated from the whole set of genotypes, both unbiased identity probability (uPI) and sibling identity probability (PIsibs) were low (uPI= 7.74e-15; PIsibs = 1.14e-05). A number of five microsatellite loci were required to achieve the conservative value of PIsibs < 0.01 (Waits, Luikart & Taberlet, 2001) (Appendix Fig. 3).

The analysis of nuclear genetic variance (PCoA) showed a lack of structuring among (i) mtDNA-assigned individuals from the WAfr and Gha lineages or (ii) countries (Fig. 3).

**Fig 3.**
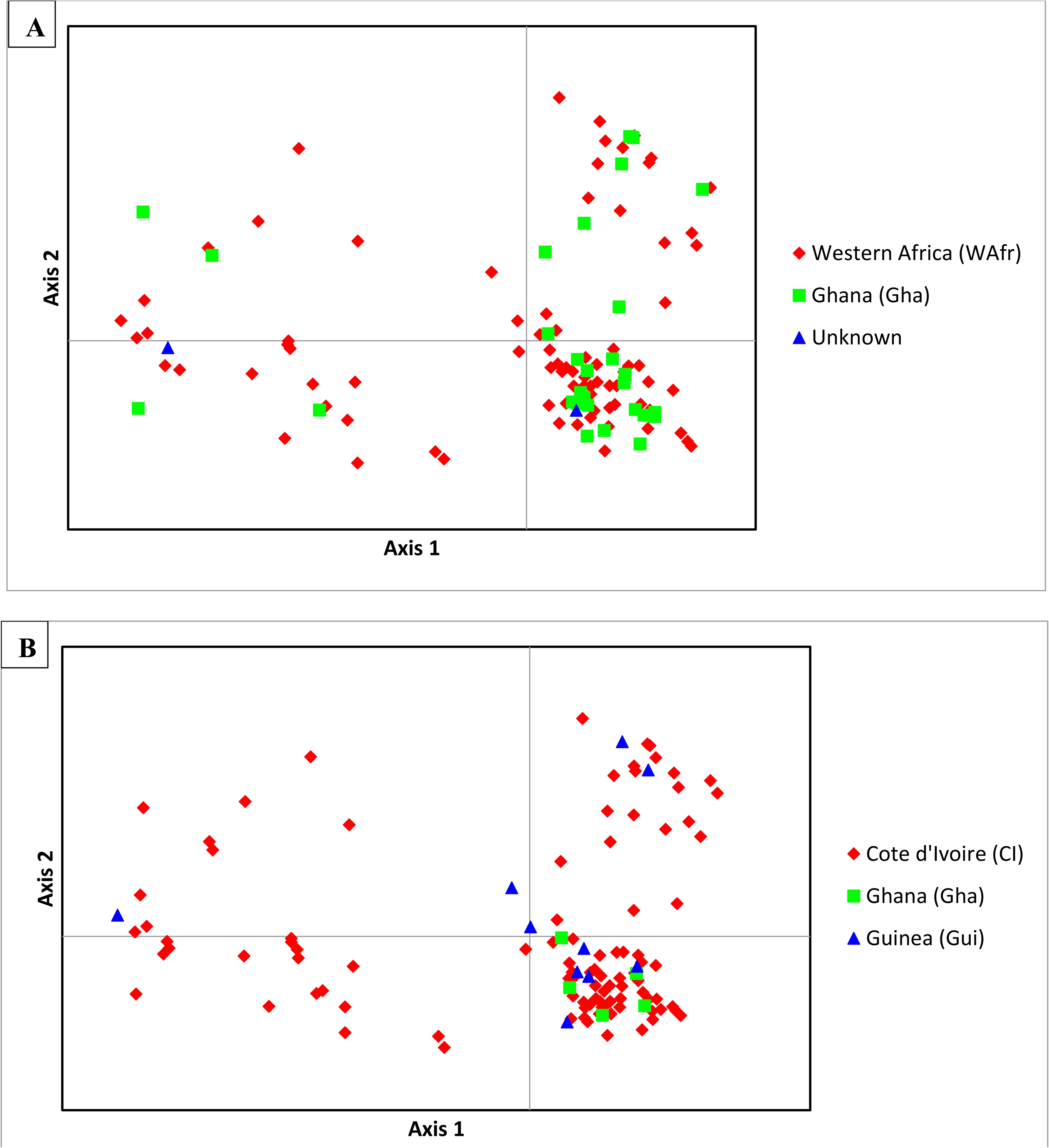
Distribution of nuclear genetic variance (PCoA) within white-bellied pangolins from West Africa according to mtDNA-delimited lineages (WAfr and Gha) (A) and countries (B). Axes 1 and 2 explain 11.35% and 8.10% of the total variation, respectively. See Appendix Figure 4 for further projections.

F_ST_ values among reference populations after the Bonferroni correction were significant for the pairs Prikro - Mont Nimba and Toumodi - Mont Nimba. Overall, F_ST_ values among populations ranged from low to moderate levels of differentiation, with a minimum value of 0.00597 (between Dagbégo and Dimbokro) and a maximum value of 0.14879 (between Prikro and Mont Nimba) (Appendix Table 5).

STRUCTURE identified three optimal clusters (Appendix Fig. 5). However, from K = 2 to K = 8 no coherent geographic structuring of populations could be observed (Appendix Figure 6).

We identified a significant IBD effect among individuals across western Africa (*r* = 0.139; p = 0.033) and among reference populations (*r* = 0.496; p = 0.025) (Fig. 4).

**Fig 4.**
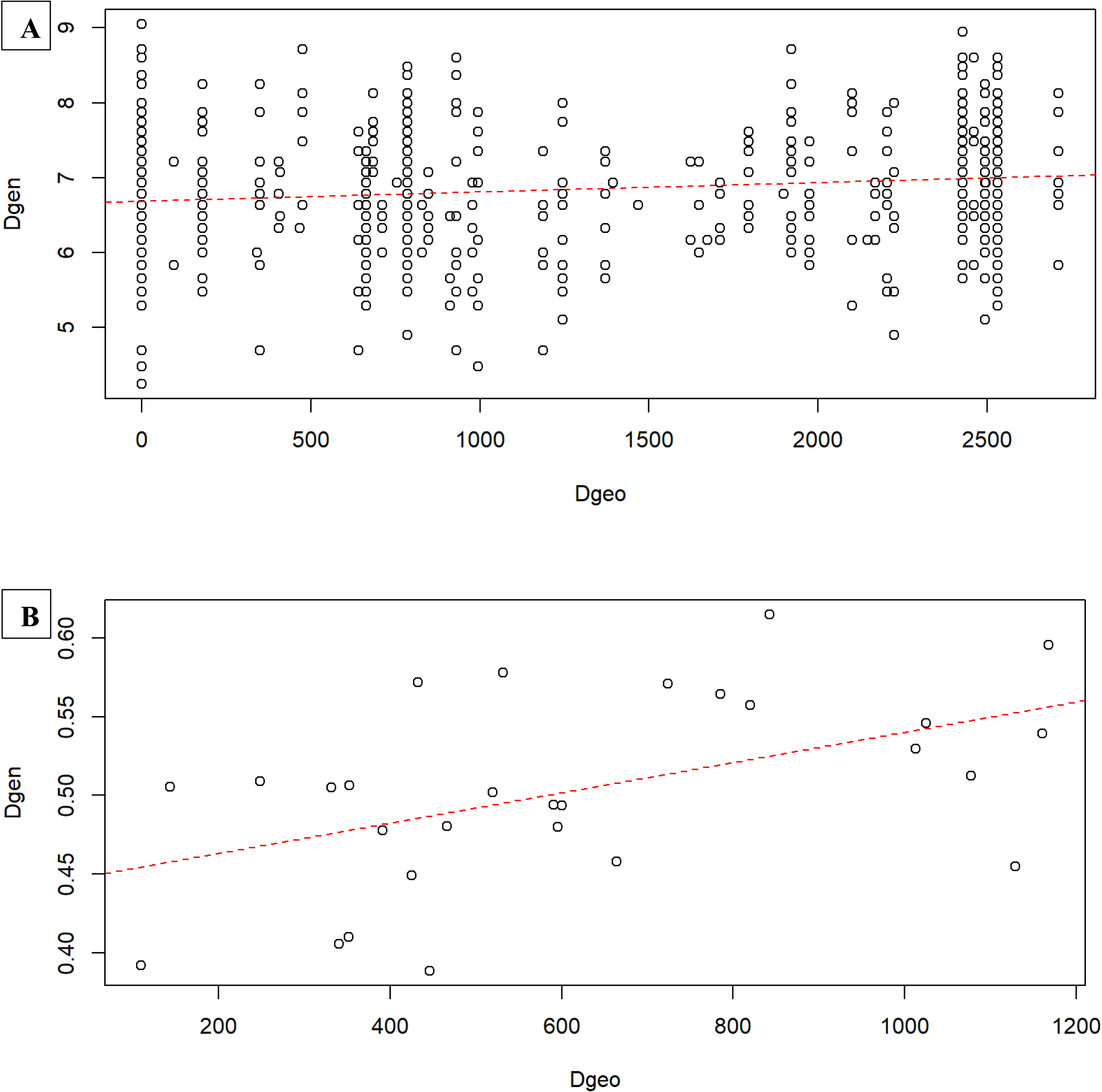
Isolation-by-distance among (A) individuals and (B) reference populations of white-bellied pangolins from West Africa, as inferred from 14 microsatellite loci. Dashed curve indicates linear regression.

VarEff showed a drastic decline in the effective population size (*Ne*) of WBP from western Africa under the three distinct mutation models (Fig. 5). Our results suggest 85-98 % reduction of *Ne*, from 10100-10900 (ancestral *Ne*) to 520–590 (contemporaneous *Ne*) individuals as harmonic means (95% CI 423-5766). Such decline was estimated to occur between 200 and 1600 generations (400-3200 ya).

**Fig 5.**
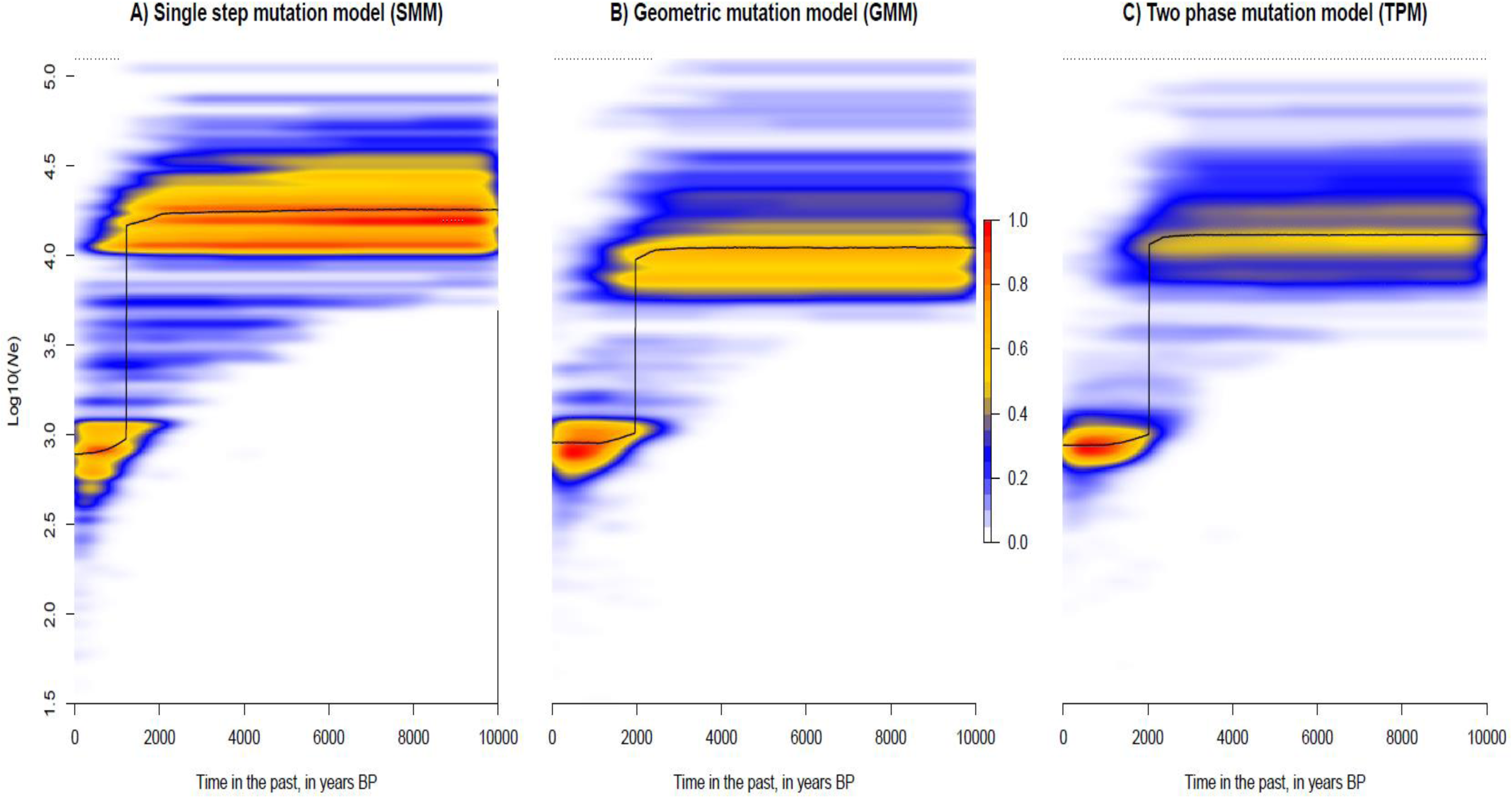
Temporal change in the effective population size of white-bellied pangolins in West Africa, as estimated from VarEff under three different mutation models. Mode (black line) and kernel density (color scale) of effective population size (*Ne*) posterior distributions are given in years BP.

## Discussion

### Lack of genetic structure between West African lineages of white-bellied pangolins

Our results showed that (i) the WAfr and Gha mtDNA lineages delineated by Gaubert et al. (2016) were geographically overlapping in eastern Côte d’Ivoire and (ii) there was a global pattern of admixture across the West African range of WBP, with no coherent genetic structuring. Our exhaustive sampling across West Africa allows us to re-delineate the ranges of WAfr and Gha lineages, the latter –previously restricted to Ghana– reaching eastern Côte d’Ivoire. Because we did not detect Gha haplotypes west of the Bandama River and WAfr haplotypes east of the Comoé River, the two rivers may have acted as barriers to the post-refugial spread of the two mtDNA lineages (see Gaubert *et al*., 2016). Although previous investigations have suggested that the two rivers could have acted as biogeographic barriers for other terrestrial vertebrates (Nicolas *et al*., 2010; Leaché *et al*., 2019), our results are based on a relatively reduced number of samples and should be considered with caution, especially since we obtained a contrasting picture from the microsatellites data (see below).

Beyond reshuffling the delimitation of the two West African mtDNA lineages of WBP, we found a general pattern of nuclear admixture across the West African species’ range, where Ghana was not differentiated from the rest of West Africa. This may be due to a high rate of gene flow between the two lineages following range expansion from Pleistocene refugia, the signature of which still being prevalent in the coalescence of mtDNA (see Gaubert *et al*., 2016). To our knowledge, it is the first time that such scenario is posited as part of the biogeography of West African mammals.

Seminal study on the genetic diversity of WBP in West Africa similarly concluded to a lack of genetic structure within the Dahomey Gap lineage (Zanvo *et al*., 2022), suggesting unexpected dispersion ability of the species. We also detected a significant IBD signature both among individuals and reference populations across West African WBP, and a few significant differentiations (F_ST_) with the westernmost population from South-East Guinean forests reinforcing the IBD pattern. Although the uneven sampling of reference populations across the study zone prevents us from concluding firmly on this pattern, it is possible that greater than expected dispersion distances are shaping the distribution of genetic diversity of WBP in West Africa. The dispersal ecology of WBP is literally unknown. However, long-range dispersal has been reported and greater than expected mobility was predicted in terrestrial species of pangolins (Van Aarde, Richardson & Pietersen, 1990; Pietersen, McKechnie & Jansen, 2014b; Ching-Min Sun *et al*., 2020). As the habitat requirement of WBP notably rely on the presence of large trees (Gaubert, 2011), long-range dispersal implies continuous forest cover across the study zone, a condition not necessarily met in West Africa due to high rates of deforestation (Aleman, Jarzyna & Staver, 2018). Further studies using tracking devices are necessary to fill the knowledge gaps on the ecology and dispersal ability of the species (Heighton & Gaubert 2021).

The lack of nuclear delimitation between the two West African lineages of WBP has serious implications on the utility of using mtDNA alone as tracer of the pangolins’ trade. Recently, mtDNA-typing following Gaubert *et al*. (2016)’s lineage delimitations was applied to pangolin scale seizures from Hong Kong (Zhang *et al*., 2020). The study concluded to the detection of 73 WAfr and 12 Gha samples from two seizures in Kenya and Nigeria. However, from our results any traced individual attributed to WAfr and Gha should be considered as originating from a wider range englobing Ghana as the easternmost possible origin, Côte d’Ivoire and Guinea (and likely the westernmost part of the species’ range). Based on the admixture pattern that we observed, we advocate for a multi-locus tracing strategy taking into account the limitations –in terms of geographic resolution– of the genetic resources available so far for WBP.

### Low genetic diversity and demographic decline in white-bellied pangolins from West Africa

The level of mitochondrial diversity was low in the Gha and WAfr lineages, coming second after the most genetically depauperate Dahomey Gap lineage. Mean effective number of alleles was slightly above what was found for the Dahomey Gap (3.099 vs. 2.491, respectively; as recalculated from Zanvo *et al*., 2022’s original dataset), whereas it was lower than in Cameroon (3.384; as recalculated from Aguillon et al. 2020’s original dataset). Our results show a global pattern of genetic pauperization in WBP from West Africa, compared to western-central and central Africa (Gaubert *et al*., 2016; Aguillon *et al*., 2020; Zanvo *et al*., 2022), although nuclear-based investigations on the species are still preliminary in ranges outside West Africa.

Although levels of inbreeding were in some populations positive and significant, values were low (≤ 0.1) so we cannot conclude that in the case of West African WBP inbreeding was one of the driving factors for low genetic diversity (contrary to the Dahomey Gap; Zanvo *et al*., 2022). We posit that admixture between the two previously isolated lineages (WAfr and Gha) could have contributed to counter-balance, at least partly, the effect of inbreeding in pangolin populations from West Africa (see Keller *et al*., 2014).

Our analyses concluded to a sharp decline in the effective population size (*Ne*) of WBP from West Africa in the recent past, c. 400 to 3200 ya. We estimated a size reduction of 85-98 %, close to the lower end of the conservative thresholds of minimum viable population size (Reed *et al*., 2003; Clabby, 2010) but slightly less drastic in amplitude than what was suggested for the Dahomey Gap lineage (Zanvo *et al*., 2022). The low genetic diversity observed in West Africa may be linked to such recent decline in *Ne* (see Charlesworth, 2009). It remains difficult to relate the period of WBP decline to a specific paleoclimatic or human-driven event, as both superimpose during the last 3200 yrs in West Africa. Since c. 12,000 yrs, a succession of abrupt periods of drought have affected the West African rainforest zone until 1300-600 ya (Hassan, 1997; Nguetsop, Servant-Vildary & Servant, 2004). Those, together with the expansion of agriculture c. 2200 ya (Ozainne *et al*., 2014), could have shaped the decline of WBP in West Africa.

Eventually, low genetic diversity and recent, sharp demographic decline may have a deleterious impact on the fitness and survival of West African WBP (see Newman & Pilson, 1997; Frankham, 2005). Additional studies based on nuclear genomic markers (SNPs) will have to be conducted to further improve our estimates of genetic diversity and demographic history of those populations.

### Potential of the genetic toolkit to trace the pangolin trade in West Africa

The mtDNA tree showed that all the samples originating from large urban bushmeat markets in Côte d’Ivoire (Abidjan) and Ghana (Kumasi, Mankessim) represented the two lineages (WAfr and Gha) endemic to the study zone. This suggests that in West Africa occur two endemic markets of WBP, one sourcing West African pangolins west of Togo and one sourcing pangolins from the Dahomey Gap, from Togo to south-western Nigeria (see Zanvo *et al*., 2022). Such demarcation is in contrast with long range trans-national trade of WBP as observed from seizures (Zhang *et al*., 2020), notably in Nigeria (Emogor *et al*., 2021). In fact, large bushmeat markets from West Africa might not represent direct hubs for the international trade of pangolins, but rather domestic hubs that source large volumes of pangolins at the national level (and from neighboring countries with close-by frontiers; Zanvo *et al*., 2022). However, denser sampling in Ghanaian urban bushmeat markets will have to be conducted to verify this hypothesis, as those markets were represented in our study by only three samples and there is a possibility that WBP from the Dahomey Gap are sold (from the neighboring Togo).

Genetic diversity indices such as the number of haplotypes and number of private alleles were strikingly high in Yopougon, the main bushmeat market from Abidjan. Such result means that Yopougon, fed by a large sourcing network (Paul & Oumar, 2011), is likely selling pangolins from a wide spectrum of locations, in Côte d’Ivoire and possibly Ghana (as Gha haplotypes were also found in the market). This is supported by the 10 different sources cited by vendors when asked about the origin of the pangolins on sale at Yopougon (Appendix Table 1), those sources being situated c. 62 - 459 km away from Abidjan. On the other hand, the high number of private alleles found in Yopougon (45 *vs.* 0-6 in reference populations) also means that we have not sampled enough reference populations across the study zone to accurately trace the pangolin trade.

Microsatellites markers were powerful enough to differentiate between individuals, and we here confirm that microsatellites genotyping is a theoretically valid approach to apply in pangolin forensic, notably on scale seizures in order to estimate the number of seized carcasses (Singh *et al*., 2020; Zanvo *et al*., 2022). We illustrated two cases where samples from a same pangolin was collected twice or thrice in Yopougon. This proves the utility of our markers to detect upstream bias in sample collection, notably when relying on third parties such a local assistants trained to assist the survey (Din Dipita *et al*., 2022). It also shows the usefulness of such markers to trace the pangolin trade, where scales and body (meat) often go through different local-to-global trade networks (Xu *et al*., 2016; Ingram *et al*., 2019; Zanvo *et al*., 2021).

Sampling of additional reference populations across the study zone, together with denser sampling within populations (see Chakraborty, 1992) will be necessary to further trace the WBP trade in West Africa.

## Conclusion

On the basis of admixture pattern, we conclude that the WAfr and Gha mtDNA lineages should be considered a single management unit (MU; Moritz, 1994; Taylor & Dizon, 1996) of WBP. We showed that this MU suffered from genetic diversity erosion and drastic decline in effective population size, and was widely sourced by at least one large urban bushmeat market in Côte d’Ivoire. It is also affected by high rates of deforestation (Norris *et al*., 2010) and a permissive national protection status, notably in Côte d’Ivoire where the species is “partially protected” and can be “hunted and captured by holders of hunting or capture permits within the limits indicated in the permit” (law n°94–442 of August 16th, 1994 modifying law n°65–255 of August 4th, 1965 relative to the protection of wildlife and the exercise of hunting). For these reasons, we consider WBP from West Africa as a MU of high conservation concern. Revision of national species status together with law enforcement and awareness campaigns should be urgently conducted, and the status of protected areas reinforced (Gonedelé Bi *et al*., 2017; Gonedelé Bi *et al*., 2019).

Our study is the first to provide comprehensive genetic evidence on the status of West African WBP. In practical terms, authorities and stakeholders in charge of sustainable wildlife management from the subregion should take advantage of this level of information to discourage the trade of pangolins. Future research on the natural history and habitat requirement of the species is urgently needed to establish a sounded, cross-national conservation strategy of WBP in the subregion. Further efforts on the genetics of the species will have to conducted, both in terms of reference sampling and development of more powerful genomic markers (e.g., SNPs), to accurately trace the WBP trade in West Africa.

## Acknowledgments

KJG and SGB received funding from the Ministère de l’Enseignement Supérieur et de la Recherche Scientifique (MESRS) of Côte d’Ivoire, as part of the “Trace-Brousse” project financed by the Franco-Ivorian Cooperation within the framework of the Debt Reduction-Development Contracts (C2Ds) managed by the Institut de Recherche pour le Développement (IRD). KJG received a PhD scholarship from “Appui à la Modernisation et à la Réforme des Universités et Grandes Ecoles de Côte d’Ivoire” (AMRUGE-CI_N°2). Field research and sampling were conducted under the authority of Ministère des Eaux et Forêts of Côte d’Ivoire, through Direction Générale des Forêts et de la Faune and Direction de la Faune et des Ressources Cynégétiques. PANGO-GO (ANR ANR-17-CE02-0001) and BUSHRISK (FCT IC&DT 02/SAICT/2017 - n° 032130) supported the lab work. PG received support from Appel à projets Actions Thématiques Muséum “Biodiversité actuelle et fossile”. Francesco Maria Angelici and Fabio Petrozzi (Italian Foundation of Vertebrate Zoology, Roma, Italy), Mac Elikem Nutsuakor (Kwame Nkrumah University of Science and Technology, Kumasi, Ghana), Brice Aymar Kouassi (Université de San-Pédro, Côte d’Ivoire) and Christophe Voisin (Muséum National d’Histoire Naturelle, Paris, France) contributed to sample collection. We thank the plateau Biologie Moléculaire et Microbiologie (B2M) at EDB – Toulouse, for teschnical facilities and support.

---

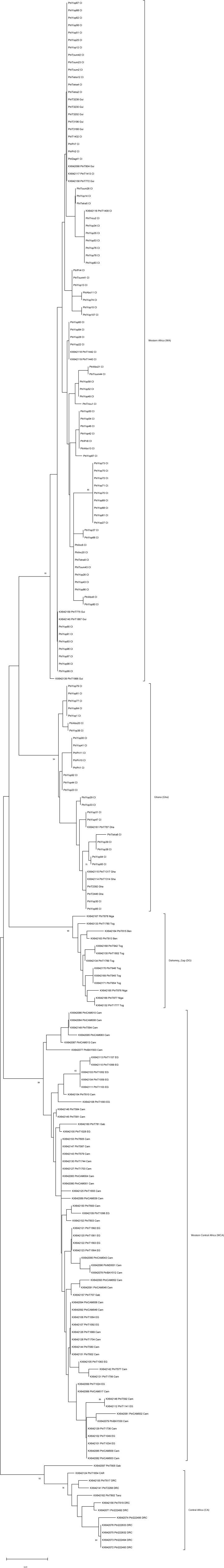

**Appendix Table 1.** Meta-data of the samples collected as part of this study, including genotypes based on 14 microsatellites markers. [Excel file]

**Appendix Table 2.**
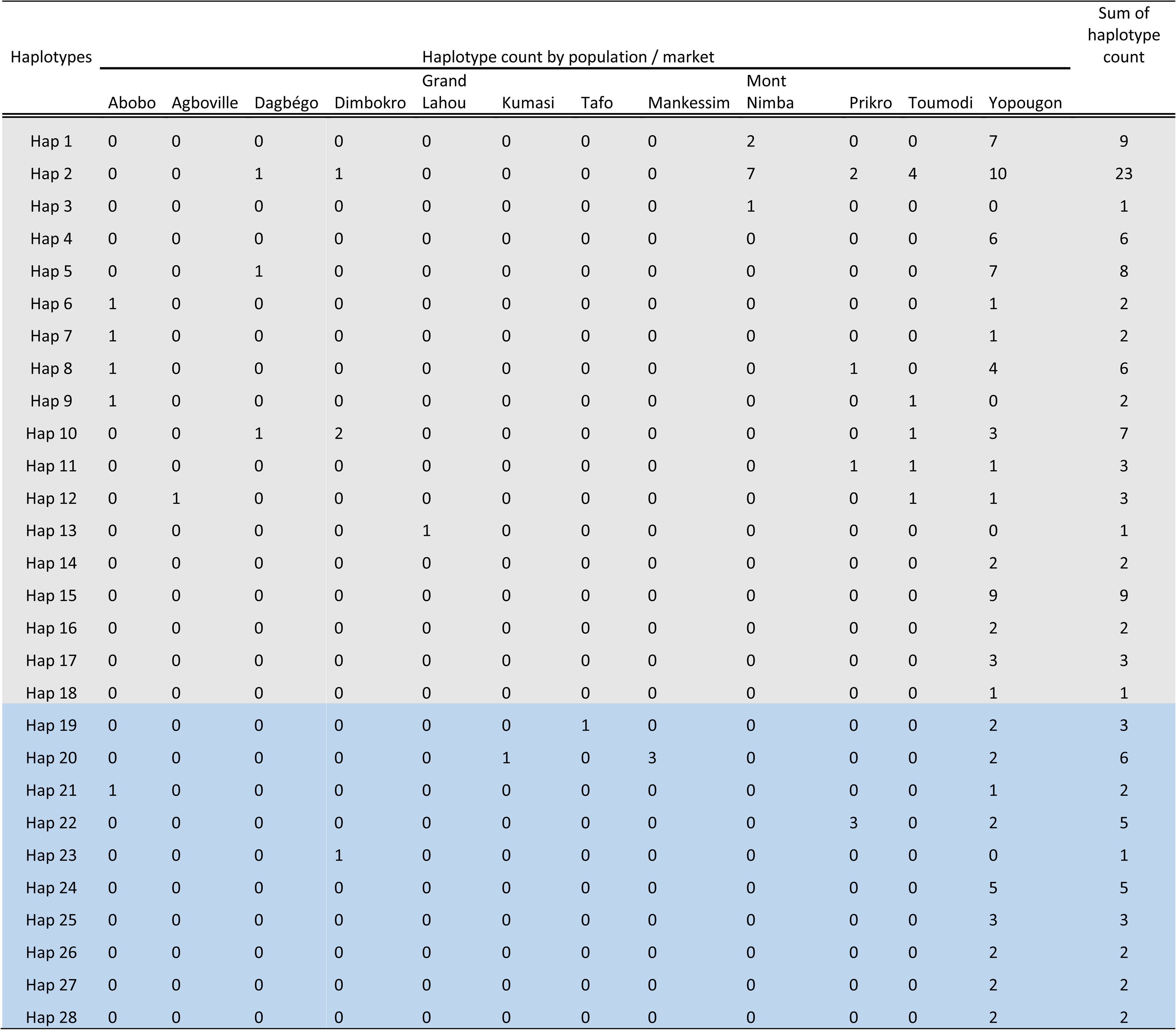
Haplotype distribution in white-bellied pangolins from West Africa. In grey, haplotypes belonging to WAfr lineage. In blue, haplotypes belonging to Gha lineage.

**Appendix Table 3.** Test of deviation from Hardy-Weinberg equilibrium per locus in the Western African lineage of white-bellied pangolins (N = 24). Significance levels assessed after Bonferroni correction. ns=not significant, (*) P<0.003.

| **Lineages** | **Locus** | **DF** | **ChiSq** | **Prob** | **Signif** |
| --- | --- | --- | --- | --- | --- |
| Western Africa (WA) | PT1453906 | 6 | 56,100 | 0,000 | * |
| Western Africa (WA) | PT739516 | 1 | 24,000 | 0,000 | * |
| Western Africa (WA) | PT34432 | 6 | 17,183 | 0,009 | ns |
| Western Africa (WA) | PT464918 | 15 | 28,995 | 0,016 | ns |
| Western Africa (WA) | PT1973508 | 91 | 118,932 | 0,026 | ns |
| Western Africa (WA) | PT796077 | 78 | 94,820 | 0,095 | ns |
| Western Africa (WA) | PT338821 | 45 | 53,411 | 0,182 | ns |
| Western Africa (WA) | PT619913 | 10 | 13,279 | 0,208 | ns |
| Western Africa (WA) | PT839522 | 55 | 62,000 | 0,241 | ns |
| Western Africa (WA) | PT2019332 | 15 | 18,024 | 0,261 | ns |
| Western Africa (WA) | PT1225378 | 6 | 7,498 | 0,277 | ns |
| Western Africa (WA) | PT308752 | 55 | 60,351 | 0,289 | ns |
| Western Africa (WA) | PT1594892 | 6 | 3,124 | 0,793 | ns |
| Western Africa (WA) | PT1669238 | 21 | 10,099 | 0,978 | ns |

**Appendix Table 4.** Distribution of null alleles per locus in the Western African lineage (N = 24).

| Locus | Null Present | Oosterhout | Chakraborty | Brookfield 1 | Brookfield 2 |
| --- | --- | --- | --- | --- | --- |
| PT1453906 | Yes | 0,2885 | 0,5152 | 0,2329 | 0,2329 |
| PT739516 | Yes | 0,1831 | 1 | 0,074 | 0,074 |
| PT619913 | Yes | 0,1586 | 0,2088 | 0,0955 | 0,0955 |
| PT796077 | No | 0,0718 | 0,074 | 0,0641 | 0,0641 |
| PT1973508 | No | -0,0519 | -0,0447 | -0,0436 | 0,1081 |
| PT839522 | No | -0,0718 | -0,0645 | -0,0645 | 0,1667 |
| PT464918 | No | -0,1026 | -0,086 | -0,0648 | 0 |
| PT34432 | No | -0,5339 | -0,2089 | -0,198 | 0 |
| PT1594892 | No | -0,106 | -0,0749 | -0,0515 | 0 |
| PT308752 | No | -0,0194 | -0,0177 | -0,0165 | 0 |
| PT1669238 | No | -0,0758 | -0,0655 | -0,0609 | 0 |
| PT1225378 | No | -0,0032 | 0,0066 | 0,0047 | 0,2101 |
| PT338821 | No | -0,0774 | -0,0682 | -0,0665 | 0,1034 |
| PT2019332 | No | 0,0019 | 0,0142 | 0,0114 | 0,2513 |

**Appendix Table 5.** Pairwise F_ST_ values as a measure of differentiation between population pairs across West Africa. Significance levels assessed after Bonferroni correction (<0.003). ns=not significant, (*) P<0.003

| Population1 | Population2 | $F_{ST}$ | p-value | Signif |
| --- | --- | --- | --- | --- |
| Prikro | Mont-nimba | 0.14879 | 0.0010 | * |
| Toumodi | Mont-nimba | 0.10032 | 0.0020 | * |
| Dimbokro | Toumodi | 0.09290 | 0.0055 | ns |
| Mont-nimba | Dimbokro | 0.07555 | 0.0059 | ns |
| Prikro | Toumodi | 0.06783 | 0.0060 | ns |
| Mont-nimba | Dagbégo | 0.07777 | 0.0060 | ns |
| Toumodi | Dagbégo | 0.03520 | 0.0104 | ns |
| Dimbokro | Prikro | 0.04253 | 0.0107 | ns |
| Dagbégo | Prikro | 0.02342 | 0.0128 | ns |
| Dagbégo | Dimbokro | 0.00597 | 0.0150 | ns |

**Appendix Figure 1.** Neighbor-Joining tree of white-bellied pangolins based on control region (CR1) sequences showing the six mitochondrial lineages clustered after Gaubert et al. (2016). [pdf file] Bootstrap values >75% are given at nodes. Scale bar represents K2P distance value.

**Appendix Fig 2.**
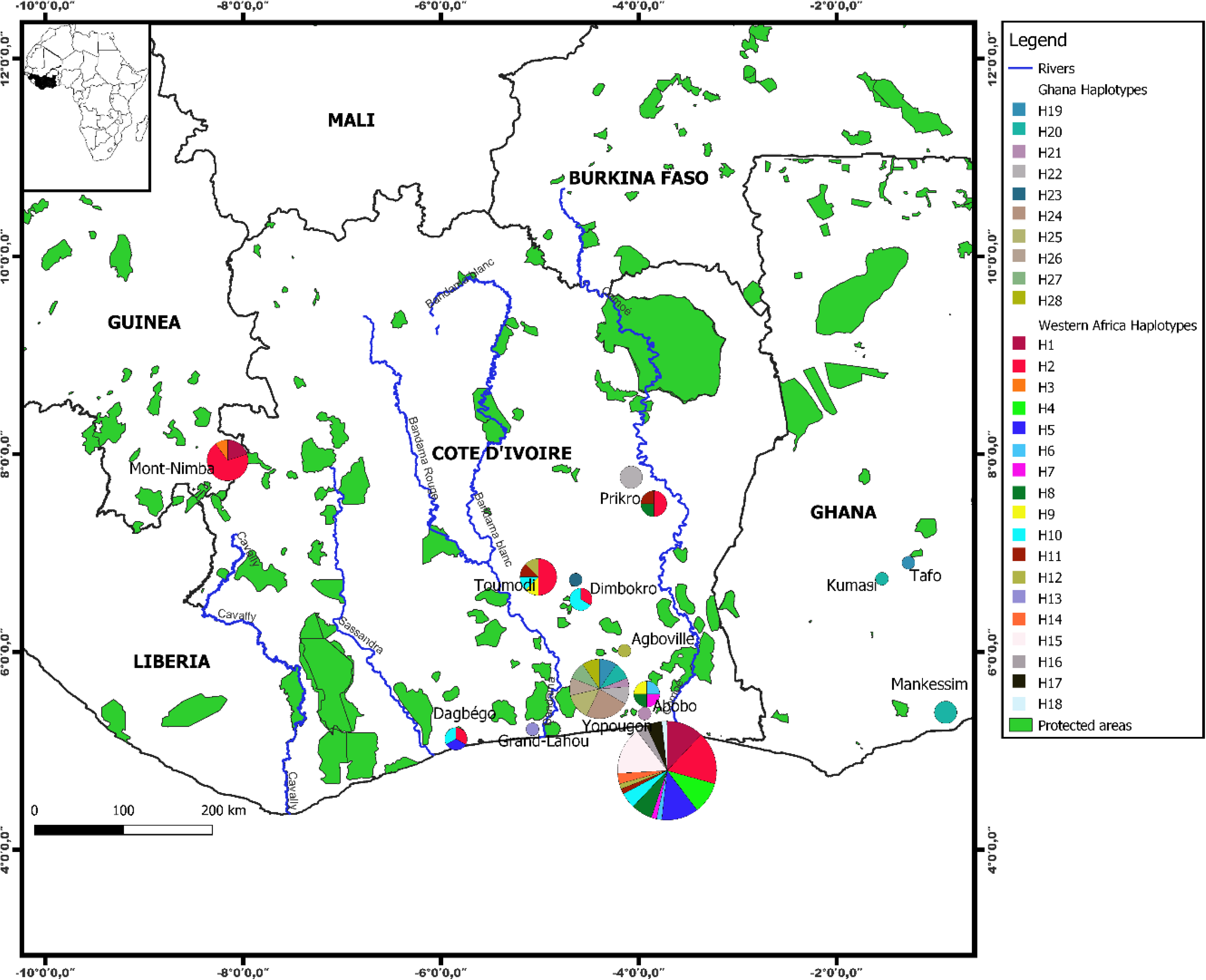
Geographic distribution of control region haplotypes across West African white-bellied pangolins. Size of pie chart is proportional to number of individuals (e.g., 1 individual in Tafo, Ghana).

**Appendix Fig 3.**
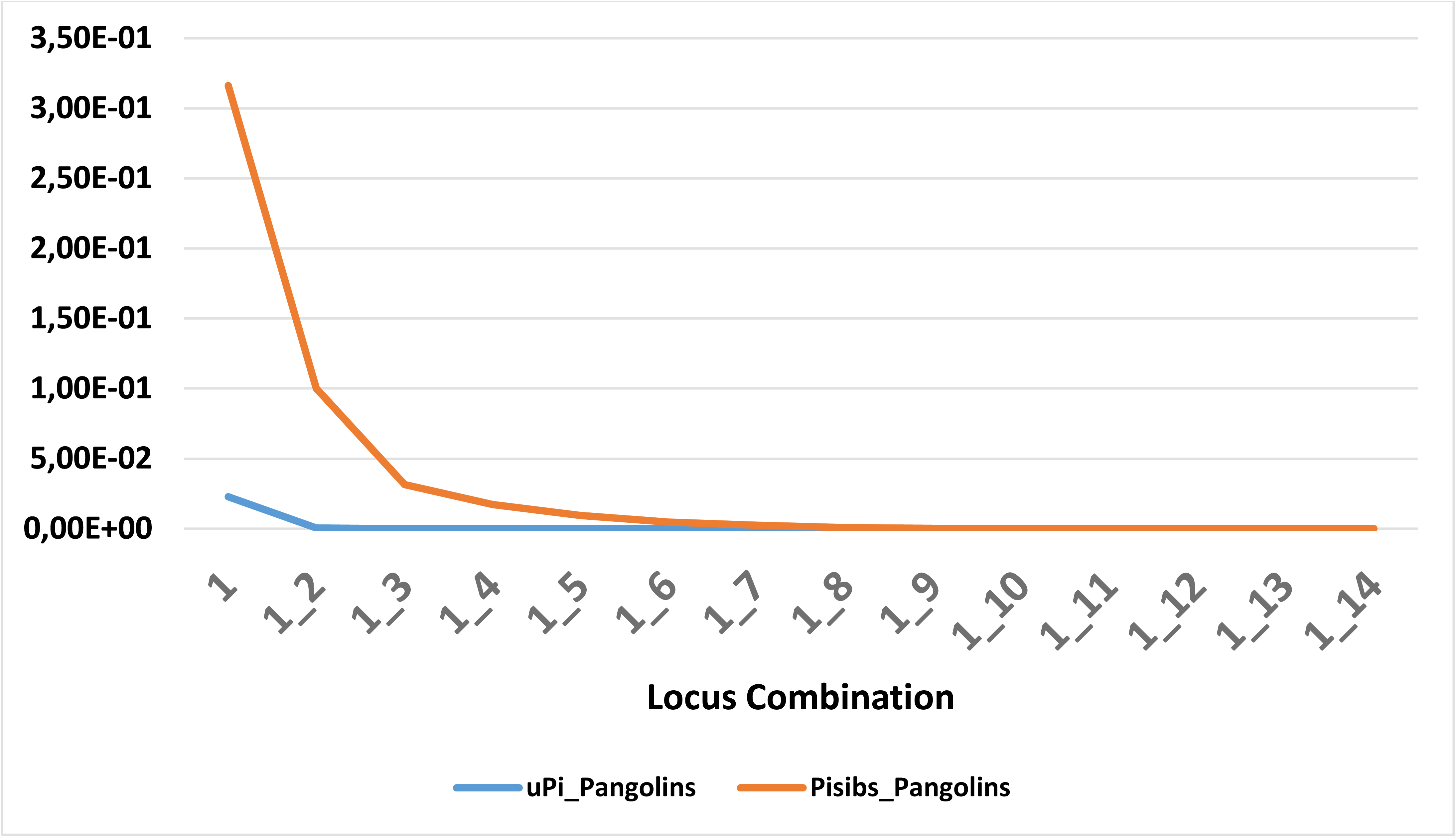
Unbiased probability of identity (uPI) and probability of identity among siblings (PIsibs) for increasing, optimized locus combinations.

**Appendix Fig 4.**
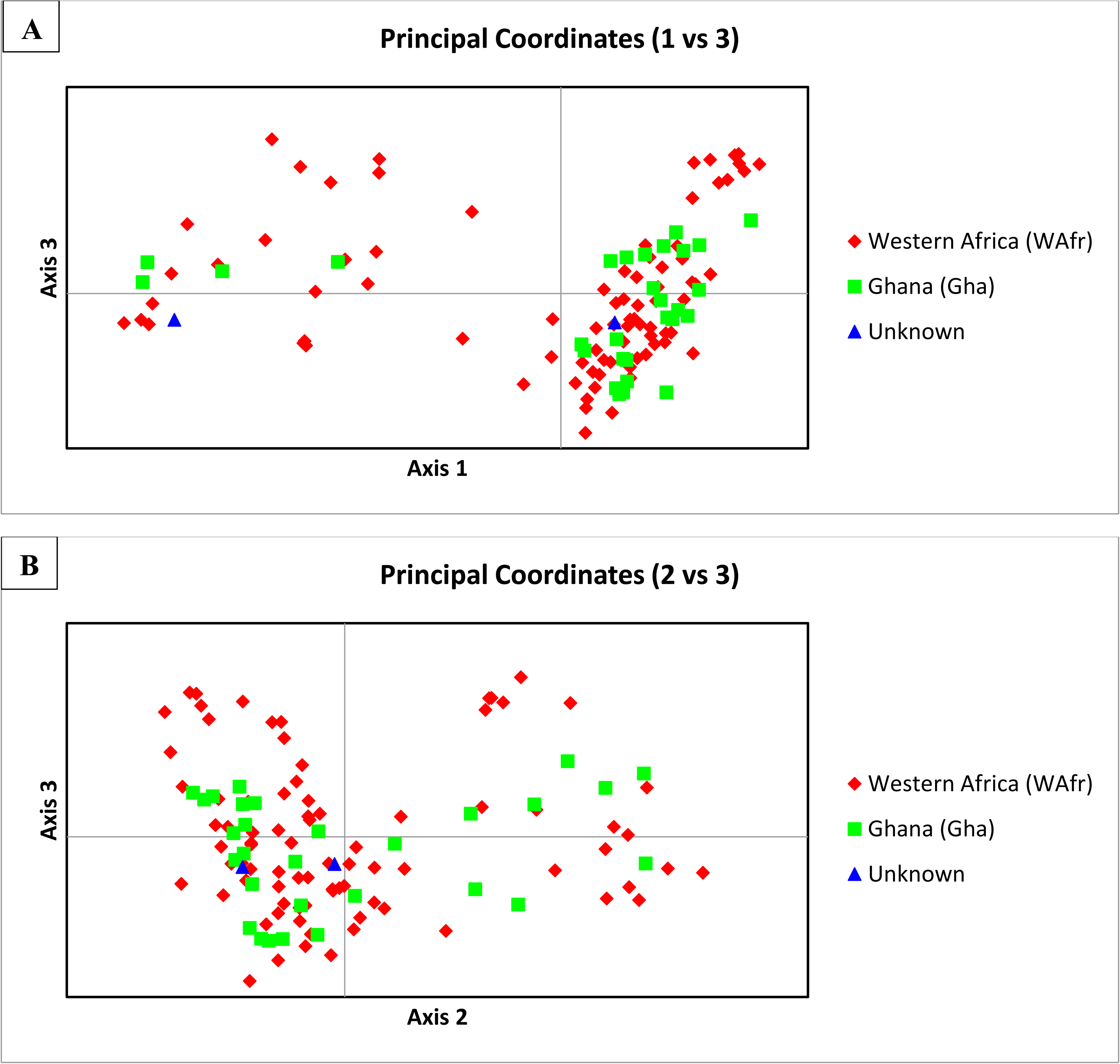
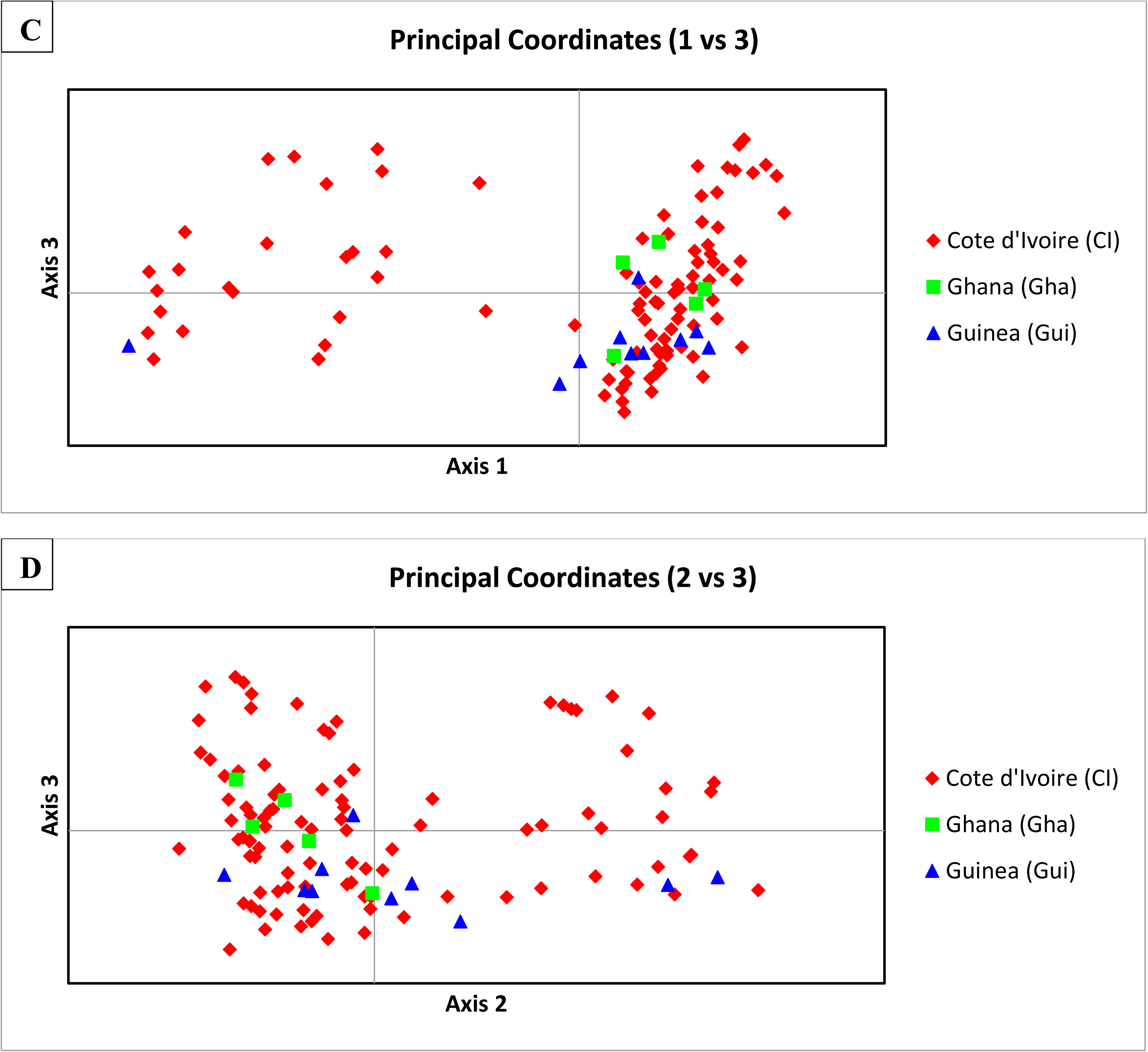
Distribution of nuclear genetic variance (PCoA) within white-bellied pangolins from West Africa according to mtDNA-delimited lineages (WAfr and Gha) (A and B) and countries (C and D). Axes 1, 2 and 3 explain 11.35%, 8.10% and 6.58% of the total variation, respectively. See Figure 3 for projections on axes 1 and 2.

**Appendix Fig 5.**
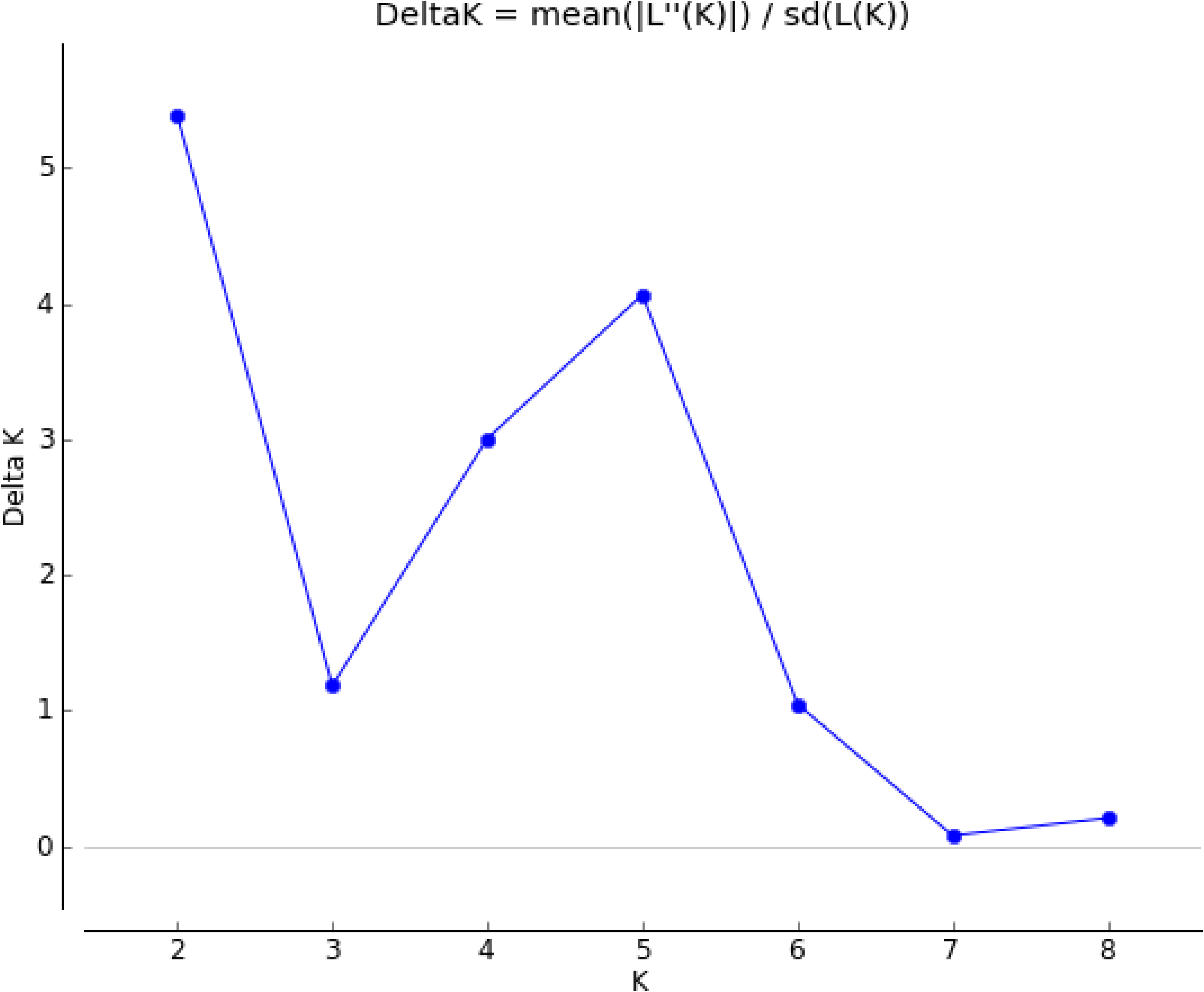
Rate of change in the log probability of the microsatellites data between successive K values (Delta K) as calculated and visualized using STRUCTURE HARVESTER.

**Appendix Fig 6.**
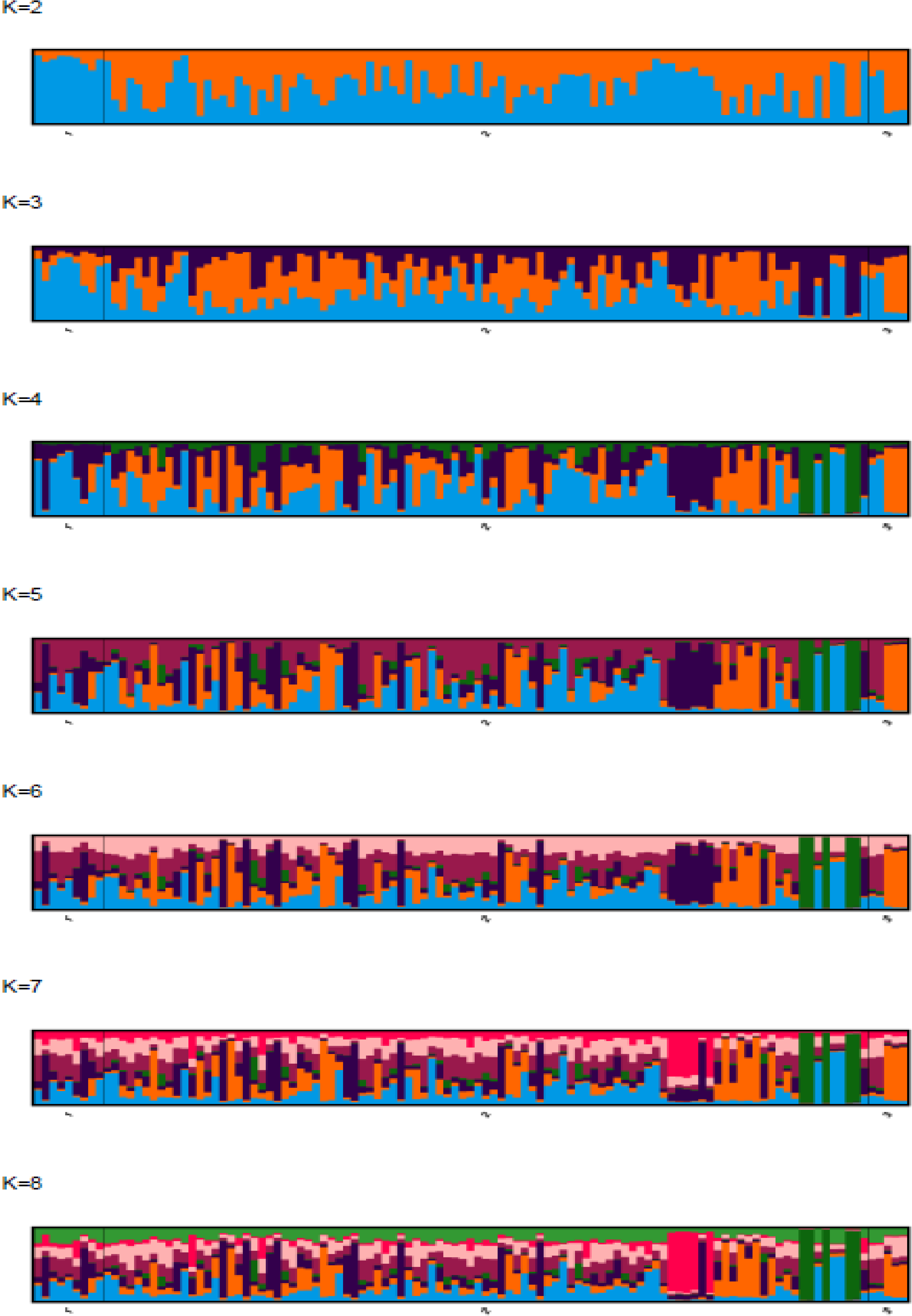
Plots of the probabilistic individual assignments inferred by STRUCTURE among white bellied pangolins from West Africa. Vertical axis represents the averaged fraction of ancestry per individual across K=2-8 populations as summarized with CLUMPAK. Populations as follows: 1- Guinea; 2- Côte d’Ivoire; 3- Ghana.

